# A semi-automated design for high-throughput Lepidoptera larvae feeding bioassays

**DOI:** 10.1101/2020.08.02.232256

**Authors:** Inoussa Sanané, Judith Legrand, Christine Dillmann, Frédéric Marion-Poll

## Abstract

Lepidopteran pests cause considerable damage to all crops over the world. As larvae are directly responsible for these damages, many research efforts are devoted to find plant cultivars which are resistant or tolerant to pest attacks. However, such studies take time, efforts and are costly, especially when one wants to not only find resistance traits but also to evaluate their heritability. We present here a high throughput approach to screen plants for resistance or chemicals for their deterrence, using a leaf-disk consumption assay, which is both suitable for large scale tests and economically affordable. To monitor larvae feeding on leaf disks placed over a layer of agar, we designed 3D models of 50 cages plates. One webcam can sample simultaneously 3 of such plates at a rate of 1 image/min, and follow individual feeding activities on each leaf disk and the movements of 150 larvae. The resulting image stacks are first processed with a custom program running under an open-source image analysis package (Icy) to measure the surface of each leaf disk over time. We further developed statistical procedures running under the R software, to analyze the time course of the feeding activities of the larvae and to compare them between treatments. As a test case, we compared how European corn borer larvae respond to quinine, considered as a bitter alkaloid for many organisms, and to NeemAzal containing azadirachtin, which is a common antifeedant against pest insects. We found that increasing doses of azadirachtin reduce and delay feeding. However, contrary to our expectation, quinine was found poorly effective at the range of concentrations tested. The 3D printed model of the cage and the camera holder, the plugins running under Icy, and the R scripts are freely available, and can be modified according to the particular needs of the users.

## Introduction

Crops are exposed to increased pressure from insect pests, partly due to climate warming which changes the distribution of pest insects [1,2]. It is also a consequence of the globalization of human activities, which allow pests to cross natural barriers and invade new ecosystems, causing massive direct and indirect economical costs [3]. Such pests often become invasive because they lack their usual natural enemies, and they face plants which may have poor defenses against them [4]. When confronted to new biotic and abiotic conditions, they may become more adaptable [5]. Invasive pests require the design of new strategies to protect cultures including swift detection and efficient measures to contain their spread [4].

To cope with pests, resistant or tolerant crop varieties can be developed but the whole process is both costly and time-consuming [6]. To facilitate the process, high-throughput laboratory assays are needed to identify plants potentially carrying resistance traits, within traditionally bred varieties or material derived from global germplasm. This can be done by growing plants either in a field or in a controlled environment and then, by measuring the impact of specific pests on the final yield or on plant organs exposed to the pest.

Leaf-disk assays are commonly used to evaluate plant resistance against insects with chewing mouthparts [7,8] by measuring the area of tissues consumed. They can be used also on mites, thrips, aphids, whiteflies, or even fungi by monitoring visual changes related to the damaged areas of the leaves [9]. While disk assays have drawbacks such as the damage inflicted to the tissue, several studies demonstrate that resistance scores obtained using this approach are comparable to those completed on detached or attached leaves [9].

Leaf-disk assays are particularly well adapted to evaluate antibiosis factors against Lepidoptera larvae feeding on leaves [10]. They are even relevant to study larvae feeding within the stem because adult females deposit their eggs on the plant surface. Therefore, first instar larvae hatching from the eggs need to graze on the leaf surface, and bore a tunnel through the leaves to reach the inner tissues. Leaf-consumption by young larvae can thus be used as a proxy to evaluate plant resistance, and to find and evaluate the effectiveness of feeding deterrents extracted from plants or of synthetic origin [10–14].

So far, very few systems have been described to perform such tests on a large number of insects at once. Most experimental setups used image analysis to measure leaf disk area consumed by few larvae either visually [7,10], or digitally [15–19]. These approaches are well suited to laboratory investigations on small scale series, but handle a limited number of repetitions and often make use of specialized and expensive hardware.

Here, we describe a fast and reliable testing protocol that can be performed using a minimum of hardware components [20], based upon measuring the consumption of leaf disks by larvae. Our approach combines three elements (i) a novel and flexible feeding bioassay whereby the consumption and movements of 150 larvae can be followed across several days with a webcam, (ii) programs to analyze the stacks of images (RoitoRoiArray and AreaTrack), written as plugins running within the free bio-image software ICY [21], and (iii) a new cutting-edge statistical approach to compare the kinetics of larval feeding [22], taking into account the diversity of behavioural responses in the insect population. As an example, we monitored the feeding behaviour of European corn borer (ECB) second instar larvae on maize leaf disks treated with different concentrations of NeemAzal and of quinine, which are considered as antifeedants for a number of insect species.

## Results

### The typology of feeding behaviours results from a novel, high-throughput feeding bioassay

We designed a high-throughput feeding bioassay to analyse the feeding behaviour of second instar larvae (L2) from European corn borer *Ostrinia nubilalis* Hübner (Lepidoptera: Crambidae) on maize leaf disks. We examined the feeding behaviour of cohorts of larvae given access to maize disk treated with different amounts of two antifeedant products. Although considerable variation exists between larvae, we showed that their behaviour can be classified into six categories (behavioural types). Differences between behavioural types relate to the time to engage into sustained consumption and to the consumption rate. The building-up of behavioural typologies relies on three innovations:

#### *i*) A custom recording system

each recording unit (Fig 1) includes three plates of 50 cages, placed over a white LED panel and monitored by a webcam mounted on a custom stand. The bottom of each plate is made of a transparent glass plate. Each cage is first filled with a 2-3 mm layer of agar on which a freshly cut leaf-disk is deposited. Then one larva is placed in each cage and a second glass plate is placed on the top to close cages. The cost of one system (including the webcam, the LED panel and the plates) is less than 400 $ US. The camera can record images at regular intervals (1 per minute) using a surveillance camera security software (VisionGS). This allowed us to register the feeding behaviours of 150 larvae, distributed into three plates, during several days. Each plate received a different treatment corresponding to different concentrations of antifeedant product. One standard computer could easily control 4 webcams that is to say it is possible to record feeding bioassay tests with 600 larvae at a time on one computer. The limiting factor is the time required to introduce insect larvae into the cages. For this reason, it is difficult for one operator alone to start more that two experiments including 300 larvae at a time, as was done here. In this paper, a single bioassay or batch corresponded to two cameras, each monitoring three plates.

**Fig 1.**
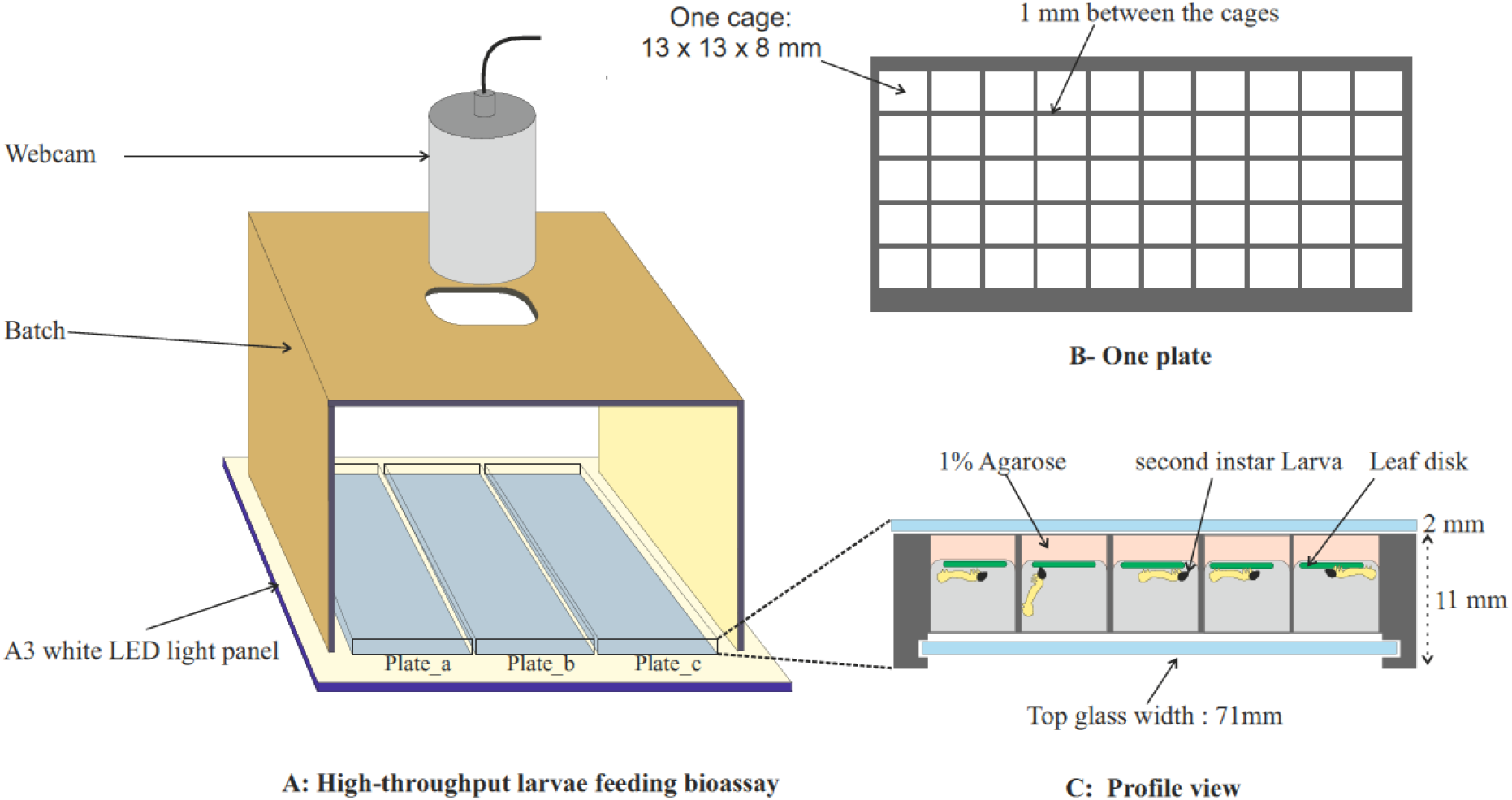
Feeding bioassay device for real-time image video recording. **A.** One feeding bioassay corresponds to one batch, that includes three 50-cages plates numbered a, b, c, and a webcam that is connected to a computer for image registration. **B.** One plate consists 50 cages, each representing an experimental unit. **C.** Profile view of one plate covered with two transparent glasses that shows larvae feeding on leaf disks during the run of an experiment.

#### *ii*) An open-source software for image analysis

the analysis of the image stacks (1920 x 1080 pixels; RGB 24 bits) is performed with 2 plugins written in Java, and running under the opensource bio-imaging software, Icy [21]. The main steps of image analysis are described in Fig 2. A first plugin, RoitoRoiArray is used to help users creating areas defining the contour of each cage, and attributing a unique ID number to each one. A second plugin, AreaTrack is used to measure the leaf area at each time point. Leaves are isolated from the background, either with a set of pre-defined image filters transforming the image into a grey image on which a simple threshold is applied, or with user-defined arrays of colours (see Methods section). The number of pixels matching these filters is monitored over time and exported as a csv file. Each csv file contains the time in lines and 50 columns, with the pixels measured in each cage at each time point.

**Fig 2.**
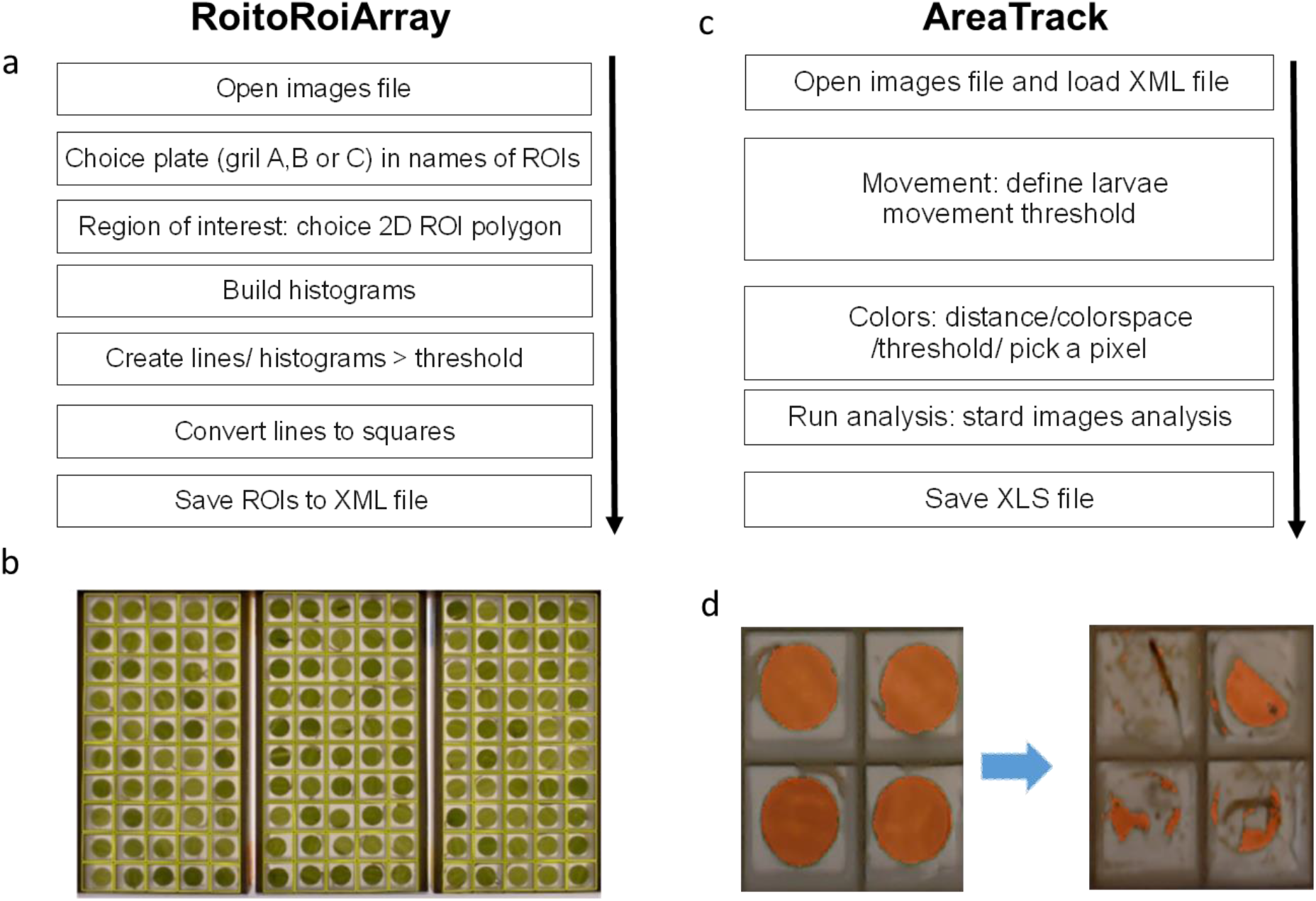
Image analysis plugins. RoitoRoiArray. (left) automatically delimits the cages. **a.** Sequence of instructions in the plugin interface that need to be run step by step by the user. **b.** image of the three plates of a batch at the beginning of the experiment. **AreaTrack** (right) measures intact leaf disks area at each time point. **c.** Sequence of instructions in the plugin interface. **d.** Examples of intact leaf areas (in orange) detected in four cages at the beginning (left) and the end (right) of the experiment.

#### *iii*) Freely-available R-scripts for data-processing and model-based data analysis

We propose a statistical method to compare treatments considering the variability of individual behaviours. One experiment may consist in several bioassays/batches. The statistical unit is a curve corresponding to the consumption of a single leaf disk measured over time (Fig 3.1). In this paper, we ran a total of four batches to compare the larvae responses to different concentrations of NeemAzal and quinine. Each batch consisted in one bioassay with two cameras moninoring a total of six plates (Table 1). Five concentrations of each of the two antifeedant product were tested. Controls consisted in a water treatment without any antifeedant. For each treatment, two plates were done and distributed among the batches (see Methods for details). Overall, we obtained 1200 consumption curves. Data analysis includes two steps. First, we define a typology of possible behaviours taking into account the whole dataset. Second, we compare the distribution of behavioural types between treatments of the same product. The different steps of the analysis, including data processing and data analysis were coded into R scripts that are freely available, along with the user manual [22].

**Fig 3.**
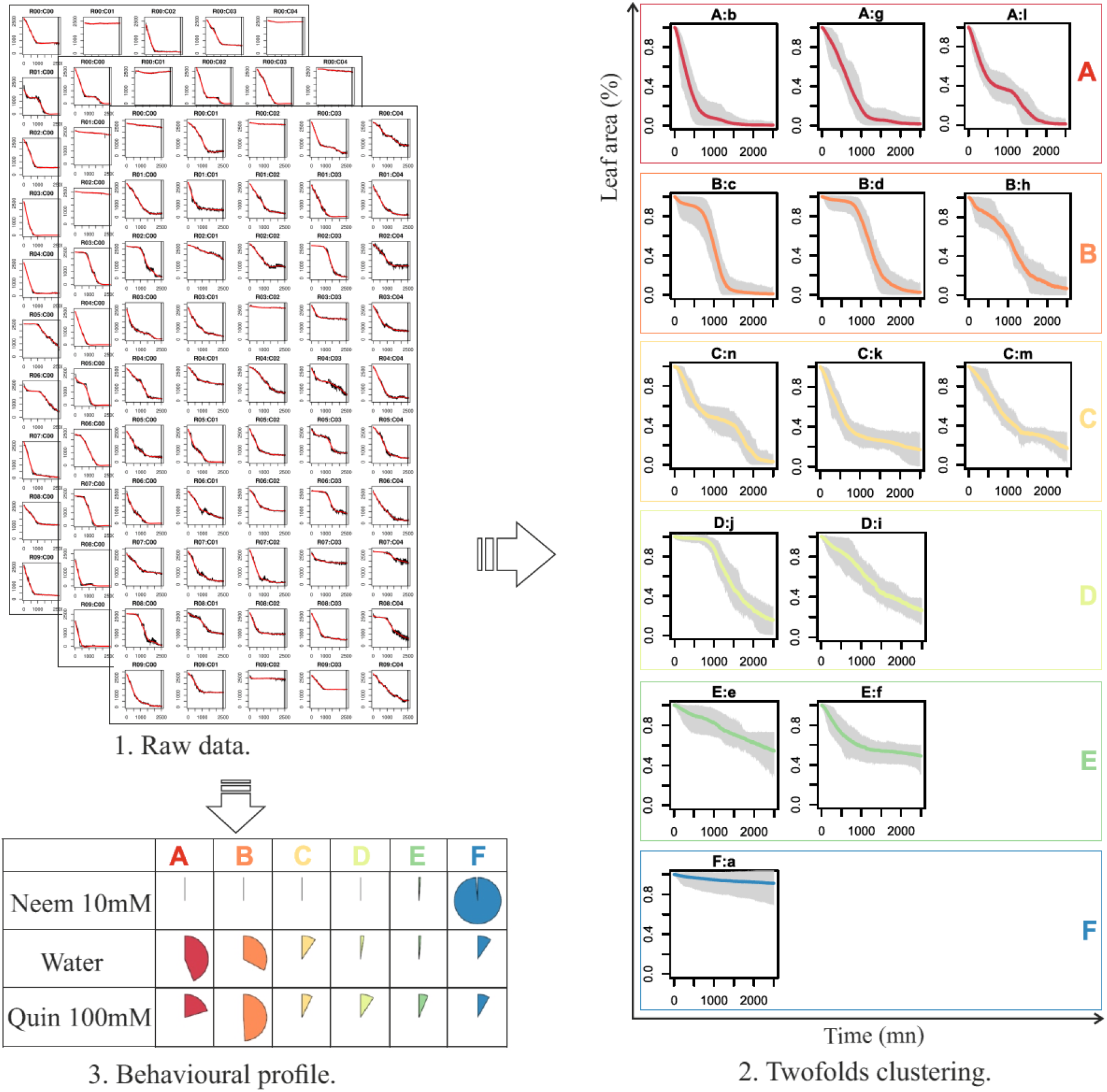
Data processing. 3.1 Raw data processing. Images are converted into (i) a pile of pdf files that allows the manual checking of the consumption curves and (ii) R data frames were each line corresponds to a cage and each column to a number of pixel at a time point. **3.2 Twofolds clustering.** First, SOTA classification results into into 14 clusters. Each cluster is represented by a vignette showing the median consumption curve (color line) and the variations around the median at each time point (grey area). Second, SOTA clusters are grouped into six types using the median value of characteristic times t20, t50, t80 and total consumption representing the clusters. At the end, the consumption curve of each cage is attributed a behavioural type, between A and F. **3.3 Distribution of behavioural types for each treatment.** As an example, distribution of behavioural types in three treatments, NeemAzal 10mM, Water (control), and Quinine 100mM are represented as pie charts. Each line sums to one.

**Table 1.** Experimental design. Each batch corresponds to an experiment performed on the same day with six different plates. One plate in a batch corresponds to one treatment. The plates are identified by letters corresponding to their position in the experimental set-up. Plates a, b, c are connected to the same camera, as well as plates d, e, f. Neem corresponds to NeemAzal treatment and Quin to Quinine treatment.

| Antifeedant | Concentration | Batch |  |  |  |
| --- | --- | --- | --- | --- | --- |
|  |  | 19-06-18 | 19-06-19 | 19-06-27 | 19-06-28 |
| Control | 00 | d | e | a | b |
| Neem (mM) | 0.01 | a | c | - | - |
|  | 0.1 | c | b | - | - |
|  | 1 | e | d | - | - |
|  | 3 | f | f | - | - |
|  | 10 | b | a | - | - |
| Quin (mM) | 0.01 | - | - | d | f |
|  | 0.1 | - | - | b | c |
|  | 1 | - | - | f | d |
|  | 10 | - | - | c | a |
|  | 100 | - | - | e | e |

#### Consumption behaviours can be classified into 6 different types

the SOTA algorithm (Self Organizing Tree Algorithm) is an unsupervised neural network associated with a binary tree topology in which terminal nodes are the resulting clusters [23]. It is particularly adapted to the classification of temporal data. We ran this method on the 1200 consumption curves and it resulted in 14 different clusters of curves having similar profiles (Fig 3.2). As shown in Figure 3.2, variations of individual behaviours around the cluster median are moderate during the first 24h. The average within-cluster coefficient of variation ranges from 3% after one hour to 25% after 16 hours. It increases up to 125% at the end of the experiment. Each cluster was further characterized by behavioural traits like the time to consume 20%, 50% or 80% of the leaf disk (noted t20, t50, t80), as well as the total consumption at the end of the experiment. Median values of clusters behavioural traits were used to group the 14 clusters into six behavioural types (Fig 3.2):

- A-type larvae immediately start to feed (low t20), consume fast (low t50, t80) and finish the leaf disk before the end of the experiment.
- B-type larvae tend to wait before consuming (high t20) but consume fast after the waiting time and generally consume all the leaf disk.
- C-type larvae consume fast in the beginning (low t20) but reduce their consumption rate through time, resulting in low t50 but high t80.
- D-type larvae show a behaviour intermediate between B and C, with high t20 and high t50.
- E-type larvae are reluctant to start feeding, showing high t20, t50 and t80, but achieving a significant consumption at the end.
- F-type larvae do not consume the leaf disk.

Each individual curve can be assigned to a single behavioural type. The comparison between treatments result in comparisons between the proportions of each behavioural type within each treatment (Fig 3.3). Figure 3.3 shows that when maize leaves are treated with water, most larvae choose the A or B feeding behaviour and consume the maize disk fast. The treatment with 100mM quinine gives almost the same results. However, when maize leaves are treated with 10mM NeemAzal, most of the larvae do not consume the leaves and are classified into the F behavioural type.

Altogether, this analysis revealed the diversity of behavioural responses of ECB larvae when feeding on maize leaf disks. Notice that even with the water control treatment, a fraction of the larvae did not consume the leaf disk.

### Bioassays confirm NeemAzal as an antifeedant for ECB larvae

After using the whole dataset to attribute a behavioural type to each individual (Fig 3), the effect of each treatment (i.e. antifeedant concentration) was analyzed separately for each antifeedant product by a multinomial regression (see Methods). Each analysis included the data of each dose of antifeedant and the controls, and resulted in the estimations of the probability distribution among behavioural types for each concentration of each product that are represented on Fig 4. A Wald test was used to test the significance of the comparisons between treatments (SI-Table 1 and SI-Table 2).

**Fig 4.**
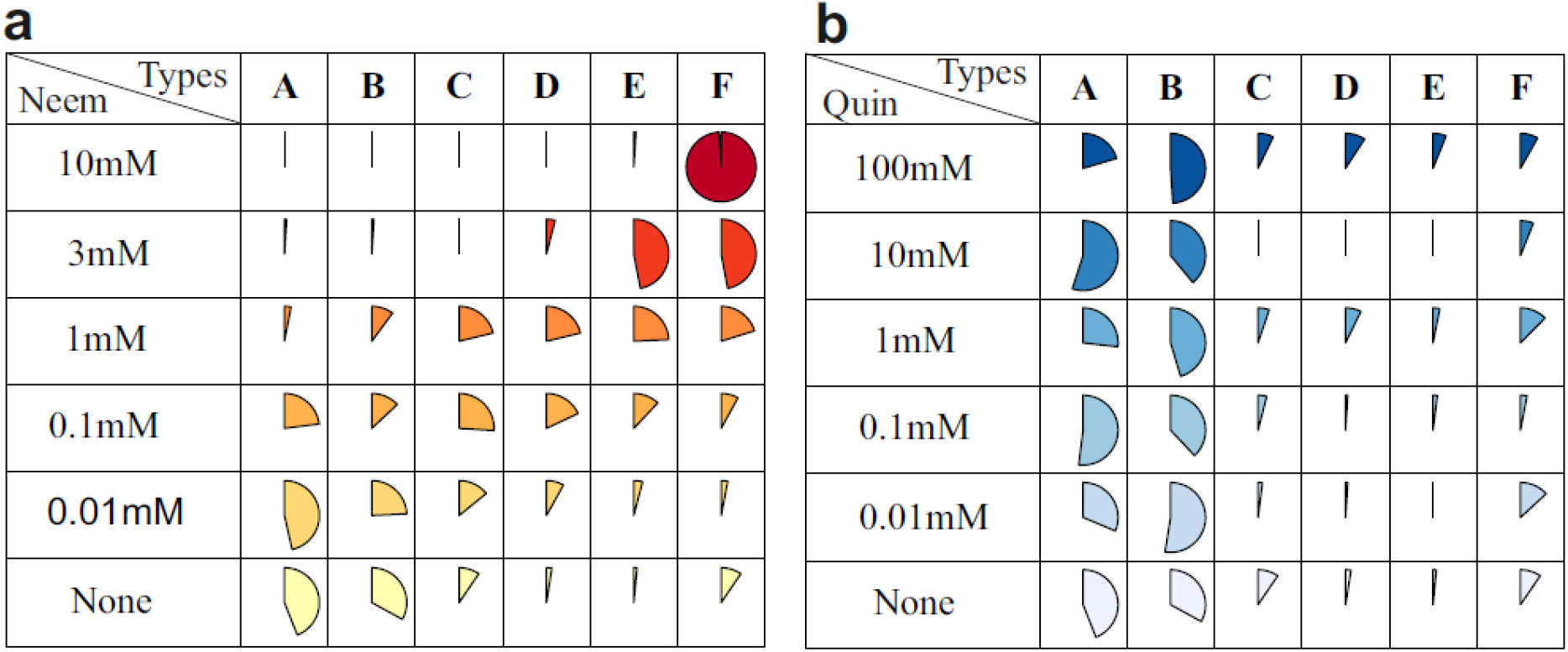
Distribution of behavioural types for the different treatments. For each treatment (row), the estimated probability of each behavioural type is represented as a pie. Each row sums to one. (a) **Neemazal treatments**. The Neemazal concentrations are indicated in the row names and represented by the red colour gradient. « None » corresponds to the control with water. (b) **Quinine treatments**. The Quinine concentrations are indicated in the row names and represented by the blue coulour gradient. « None » corresponds to the control with water.

In the NeemAzal bioassays, an increasing concentration of the product is clearly associated with a decreasing proportion of A and B behavioural types, and an increasing proportion of E and F behavioural types, while intermediate concentrations correspond to a higher proportion of C, D, E behavioural types (Fig 4a). Without antifeedant, the high proportion of A and B behavioural types correspond to larvae that immediately start consuming (A-type) or wait a short time (B-type) but then fastly consume the whole leaf disk. For intermediate concentrations of the antifeedant, a large fraction of the larvae start consuming at a lower rate (C-type) or wait and consume slowly (D-type). They generally don’t finish the leaf disk. Finally, with the highest NeemAzal concentration (10mM) 98% of the larvae are classified into the F behavioural type, that correspond to the absence of consumption.

These results are confirmed by contrast estimates relative to behavioural type E (SI-Table 1). A positive value of the contrast between treatments i and i’ means that the proportion of behavioural type w (w being A, B, C, D or F) relative to behavioural type E is greater in treatment i than in treatment i’. All contrasts between 10 mM NeemAzal and lower concentrations are significantly negative for behavioural types A, C, D, and significantly positive for behavioural type F, indicating a strong deficit of behavioural types A, C, D and a strong excess of behavioural type F in the 10mM NeemAzal treatment. Notice that for the B behavioural type, the comparisons lead to negative but non-significant contrasts. This may indicate a lack of power of the experimental design. NeemAzal treatments 0.1mM and 0.01mM are generally non-significantly different from the control, except for behavioural types A, B C. The negative value of the significant contrasts always indicate an excess of behavioural types A, C, and C in the control. Altogether, the analyses indicate that ECB larvae detect NeemAzal as an antifeedant, and change their feeding behaviour according to the concentration of the product.

In the quinine bioassay, we hardly saw any effect of the antifeedent concentration on the distribution of behavioural types, except for a slight excess of feeding types C, D, E in the 100mM treatment as compared to 10 mM (Fig 4b). This result is comforted by the contrasts that were hardly significant (SI-table 2). Even when leaf disks were treated with 100 mM quinine, the proportion of larvae showing an A/B feeding behaviour was still elevated. This indicates that quinine, when diluted with water, has no effect on larvae feeding behaviour and that most larvae consume the leaf disk entirely although a fraction do not start to consume immediately.

## Discussion

Insect feeding bioassays are of importance for plant health, and particularly plant breeding programs. They allow to evaluate plant resistance against insects with chewing mouthparts by measuring the area of tissues consumed. Here, we propose a new fast and reliable feeding bioassay based upon measuring the consumption of leaf disks by insect larvae. We set-up a high-throughput semi-automated design able to follow the feeding activities and movements of 150 larvae at a time using a single webcam. This system was built with on-the-shelf readily available elements and can be duplicated and adapted to different situations [24]. While similar systems were proposed in the past, using videographic analyses to measure leaf consumption from a limited number of larvae, this is the first time such a system is demonstrated to run on such a large sample of insects. Such performances are made possible by the current evolution of the technology and because we chose to sample images at a relatively low frequency rate. Previously described approaches using videographic analyses were combining the measure of leaf areas with feeding phases and movements of the larvae, using movies acquired at 30 images per second from a single larva [19] or from a group of twelve larvae [17]. Our approach allows to monitor 150 larvae at a time. Actually, the limiting factor is the time to prepare the setups, and more specifically, the time to collect the larvae and to introduce them into individual cages. However, one operator can start two different batches a day, using two experimental devices. For example, the whole NeemAzal experiment (600 larvae) presented here was realized in two days, using four feeding bioassays devices in parallel.

One drawback of this method, is that the feeding activities are measured on excised tissue. This has three consequences. First, the excised tissue is deprived of its normal input of nutriments and it is prone to desiccation, changing its physiology and shape. Maintaining the moisture of leaf disks punched on the plant can limit desiccation during the experiment [16,19,25]. Second, the leaf disk characteristics changed over the course of the experiment, their colour sometimes turning yellowish or becoming paler. This is why we fill the cages with 1% agarose the day before the experiment [25], and sandwich them between two glass plates as shown in Fig 1. Lastly, leaf disks are mechanically excised, thus wounding the tissues, especially along the border of the disk. As reported in [9], several studies found no differences between detached/attached leaves and leaf disc assays. Nevertheless, the results obtained with leaf disks ought to be considered as a proxy precisely because of these limitations. Notice that the use of excised leaf disk allows a physical separation between the production of the plants and the location where the pest insects are held, and therefore minimizes the risks of contamination of the plant production site.

As a test case, we analysed here how European corn borer larvae modulate their feeding activities when leaf disks are treated with one potential antifeedant, either NeemAzal, which contains azadirachtin, or quinine, which an alkaloid detected as bitter by many organisms including humans. Azadirachtin is a triterpenoid, and a complex molecule, which is often considered as a universal antifeedant for insects [26,27]. NeemAzal is a commercial extract containing mainly azadirachtin and it was previously shown that it is indeed antifeedant for ECB larvae [10,15], showing both antifeedant and insecticide activities [10,26–28]. We found here that NeemAzal has a strong impact on the distribution of consumption curves (Fig 4a). As a second test compound, we tested the alkaloid quinine, which is found in the bark of *Cinchona* and *Remijia* trees [29]. This compound is used as an antimalarial agent and is used at low doses for its bitter taste to humans. It is an antifeedant for different animals, including *Drosophila*[30], and several Lepidoptera [31–34]. Interestingly, ECB larvae did not respond to quinine in our tests. Scattered observations in the literature reported the absence of bitterness of quinine in another Lepidoptera [35] and mantids [36]. While this observation is new for *O. nubilalis*, it remains to be tested whether the lack of response to quinine is species-specific. Actually, immature insect stages may exhibit different taste responses as shown in one Diptera [37] and one Lepidoptera [38]. This lack of sensitivity could concern only the specific larval stage (L2) tested here.

Our approach introduces original statistical analyses which are new for this field. We designed a rigorous statistical approach to compare the time course of feeding activities. Classical consumption tests usually rely upon comparing feeding after fixed interval(s) of time [10,11,16,39–43]. With the approach developed here, we can characterize properly the different feeding strategies adopted by several insects confronted to the same situation. While the classical approach and our approach may globally obtain comparable results, we will be able to better characterize whether plant resistance factors or externally applied chemicals have an immediate sensory effect or if they are the consequence of post-ingestive effects.

Lastly, the system described here, with its cutting edge methodology, its affordability and its possibility to test large number of insects opens up new possibilities to evaluate the variability of feeding behaviours within populations of insects.

## Materials and Methods

### Insect rearing

*Ostrinia nubilalis* Hbn. eggs were obtained from Bioline AgroSciences (France). Eclosed larvae were maintained in Petri dishes on an artificial diet (1.32 l water, 27 g agar powder, 224 g corn flour, 60 g dried yeast, 56 g wheat germ, 12 g L-ascorbic acid, 4 g vitamin mixture and minerals (Réf.0155200), 0.8 g chlortetracycline, 2 g hydroxybenzoic acid methyl, 1.6 g sorbic acid and 4 g benzoic acid), under 16:8 (light: dark) photoperiod at 70% humidity and at 26°C. Second instar larvae (10 days old) were used for the feeding bioassays.

### Maize plants

Seeds from the maize inbred line MBS847_NLN14 were obtained from Saclay’s Divergent Selection Experiments [44]. Seeds were pre-germinated in Sprouting Trays before being transferred into individual pots (4 l) containing Jiffy® premium substrate. Plants were grown in a greenhouse at 16 h light: 8 h darkness with the temperature between 21-24°C and 70% humidity. Leaf disks of 1 cm diameter were taken from mature leaves of the same leaf rank (rank 8). To obtain samples of 50 leaf disks, approximately 10 leaf disks were sampled on five different plants at the same developmental stage, so that the experiment is not destructive.

### Experimental system

Each plate was designed as a plate of 10 x 5 square cages of (13 x 13 x 11 mm) separated by 1 mm walls [24]. The bottom of the plate was designed with two grooves to slide in a glass plate cut to the corresponding dimensions. The plate was first half-filled with 1% agar solution to maintain leaf disk moisture during the experiment [25]. One leaf disk and one larva (10 days old) was added in each cage [45]. The top of the plate was then covered with a second glass plate and maintained in place with two rubber bands, to prevent larvae to escape (Fig 1). The webcam was inserted into a stand made from MDF (Medium Density Fiberboard) cut by laser cutting (Fig 1). This stand enclosed an area fitting three plates, and ensured that images were taken at the same distance and under the same angle. The plates and the stand were placed on top of an A3 white LED light panel (white 4000K; Display Concept, Brussels, Belgium), lying flat on a table. In this paper, one bioassay consisted in setting-up two experimental systems at a time, with simultaneously monitoring six plates with two webcams.

Instructions for the making-up of the experimental system are deposited on a dataverse [24] and are freely available. Plates were designed with AutoCAD software with a 3D printer (Ultimaker 2+) using white PLA (polylactic acid) filament (Ref.RS Stock No. 134-8192), at a resolution of 0,6 mm [24].

### Experimental design

Leaf disks were deposited on the 50-cages plates onto 1% agarose gel immediately after sampling to avoid dehydration. The bioassays consisted in testing the leaf consumption by the larvae in the absence or in the presence of an antifeedant. Two products at different concentrations were tested: an extract from neem seeds, NeemAzal (AMM n° 2140090 *) that contains azadirachtin, and quinine hydrochloride (Sigma Aldrich; CAS number: 6119-47-7). Leaf disks were sprayed with 10 μl (as spraying volume) of the substance with distilled water as solvent and kept 15 to 20 min at ambient temperature to let the solution dry before starting the experiment. This spraying has been done with repeater sprayer (Repeater ® E3/E3x) on one side of leaf disk. Five concentrations were used for either NeemAzal and quinine according to experimental design (Table 1). One larva (second instar) was deposited in each cage. Plates were placed in the monitoring device and larvae were allowed to feed for 48 hours at ambient temperature. A treatment (i.e. one specific concentration of the product) was tested in two 50- cages plates with 50 leaf-disks treated with the same concentration of the product. For each product, we conducted two bioassays/batches that started on two consecutive days (Table 1). In each batch, treatments were randomly assigned to the plates. As we tested two antifeedant products, this experimental design led to the acquisition of 1200 *O. nubilalis* larvae feeding curves (2 products * 6 treatments * 2 replicates * 50 leaf disks).

### Image acquisition and analysis

#### Image acquisition

Webcams (Logitech Pro C970) were connected to the USB port of a PC computer, with sufficient disk memory to store the images. Notice that one standard computer could easily control the 4 webcams, although we only used two webcams here. Time-lapse images were taken every minute by a video surveillance software (VisionGS BE 3.1.0.4; http://www.visiongs.com) as 1920 × 1080 pixels jpg files, named with a time stamp.

#### Image analysis

Image analysis of the stacks was performed semi-automatically, with the help of custom programs working within the bio-imaging open-source program ICY [21]. In order to follow the consumption of each leaf-disk, regions of interest (ROIs) must first be defined to locate each cage or disk. This is done semi-automatically with a plugin called RoitoRoiArray (http://icy.bioimageanalysis.org/plugin/roitoarray/). The plugin automatically analyses the image stacks and makes a first proposal for the ROIs, that need to be refined by the user. Then, the plugin generates arrays of ROIs, gives them a unique ID and saves them into an XML file (Fig 2a&b). A second plugin, Areatrack (http://icy.bioimageanalysis.org/plugin/areatrack/), allows users to load stacks of images, browse or display them as a time-lapse movie, and analyses them (Fig 2c&d). After loading the ROIs from the previous file, each image of the stack is filtered and thresholded to keep only the pixels corresponding to the leaf areas. Movements of the larvae can also be detected by performing image to image subtraction followed by thresholding. These measures can then be displayed or saved as a CSV file. As considerable variation of lightning and colour of the leaf disks occurred, two detection strategies were implemented. The first approach was to transform the RGB image by transforming it into a grey plane and applying a unique threshold. The filters implemented are: R, G, B, 2R-(G+B), 2G-(R+B), 2B-(R+G), (R+G+B)/3, or converting the image into the HSB space and extracting one component H (HSB), S(HSB) and B (HSB). Although this approach is satisfactory in most cases, some experiments could not be analysed this way. We introduced another approach, consisting in letting the user point at dots on the screen to build a matrix of colours. These colours were stored either in the RGB, HSV or H1H2H3 space. The user could threshold the image using a distance around these colours, L1 or L2. The distance L1 between colour c1 and c2 is defined in the RGB space as distance

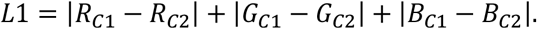

Distance L2 is defined as distance

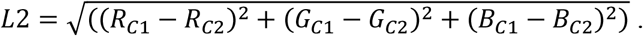

Both approaches let the user display the thresholded image as an overlay over the original images and thus choose the best adapted threshold. As these experiments generate huge amount of data, additional procedures were implemented behind the screen to fetch data from the disk in advance, and also to explore and analyse only a fraction of the data, for example one image under 10 or under 60. Once the detection was done, by analysing all images of the stack or a subset of them, the resulting data were exported as an excel file, including three type of measures: the raw data (i.e. the number of pixels over the threshold in each cage), a running average and a running median of the raw data using a user-defined span (usually n=10). While this analysis can be run on the three plates at once, it was found more convenient to perform the analysis plate by plate, so that the measures performed on each 50-cages plate were saved into one separate file. Although recordings were performed on longer time periods, this paper reports analyses made over a 48h period, corresponding to stacks of 2,880 images per plate.

### Data analysis

Series of R scripts were written for data processing and statistical analysis. The scripts, along with the raw data and a documentation were deposited in a dataverse [22] and are freely available.

#### Metadata

raw files were stored using lab storage facilities on specific directories. Directories names contain information about the date of the experiment and the camera number. Csv filenames contained all the information about the experiment: date, camera ID, plate ID, insect population, plant genotype, plant environment, plant coordinates, substance name, substance concentration. After image analysis, all csv files to be analysed together were copied under a single sub-directory

#### Standardization

first, pdf images of leaf consumption through time are produced. Each pdf file corresponds to one plate and contains 50 plots, one for each cage. Cages that produced abnormal plots, identified by their row and column coordinates, were manually removed from the analysis. Then, we standardized the data by dividing the leaf area by the total leaf area at time *t0.* Hence, the basic measure became the fraction of intact leaf disk from each cage at each time-point. We defined *tmax=2500mn* (40 hours), and only retained the data taken before *tmax.*

#### Data clustering

the whole dataset, comprising the experiments performed with NeemAzal quinine and the control (four batches, 24 plates, 1200 cages corresponding to 1200 consumption curves), was used to run unsupervised clustering algorithm SOTA [23] on the individual curves in order to obtain 14 clusters. The total number of clusters was empirically chosen to avoid having clusters containing only a couple of curves and proved to be robust over the different experiments.

#### Typology of feeding behaviour

each curve was characterized by the time after which 20%, 50% and 80% of the leaf disk was consumed, respectively noted t20, t50, t80, and the fraction of the leaf disk consumed at *tmax.* When less than 20%, 50% or 80% of the leaf disk was consumed at *tmax*, the corresponding variable was given the value of *tmax.* Then, each cluster was characterized by the median t20, t50, t80 and total fraction consumed, using the values of the curves belonging to the cluster (Fig 3.2). We then arranged the 14 clusters into 6 groups based on their median values for t20, t50, t80 and total fraction consumed using the K-means algorithm [45]. Each group corresponded to a different feeding behaviour named by a letter from A to F (Fig 3.2). All the curves belonging to the same behavioural group were assigned the same letter and are referred as “behavioural type” below and in the results section

#### Data transformation

at the end of the analysis, each cage l, corresponding to treatment i, batch j and plate k was characterized by the feeding behavioural type (A to F) of the group to which it belongs. Hence, the observations of a single cage can be summarized into a vector of zeros and ones 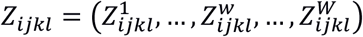, (1) where w represents one feeding behaviour, with w in {A, B, C, D, E, F}, and 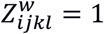 if the observed feeding behaviour is w, and 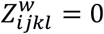 if else. For a given cage subscripted by *ijkl*, 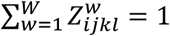

#### Statistical analysis

a separate analysis was conducted for each product (NeemAzal and Quinine). Z_ijkl_ is the result of one multinomial sampling in

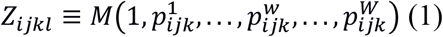

where 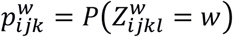 is the probability that a single observation falls into the feeding behaviour *w*. We used the logistic multinomial regression [46] to estimate the probabilities. For a given

For a given product, the full model writes

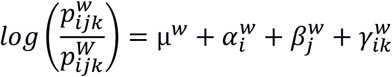

where 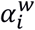 is the treatment effect, 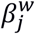 is the batch effect, and 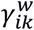 is the interaction between plate and treatment effects.

The experimental setting was highly imbalanced. For example, only control plates from the two quinine batches were added to the NeemAzal experiment, so that it was not possible to test for the batch effect. Similarly, because we used only two plates for each treatment (except for the controls), interactions between plates and treatments cannot be estimated. We therefore used the submodel (2) to infer the effects of the treatments, that writes:

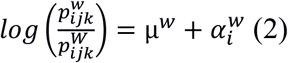

where μ^w^ is the average proportion of the feeding behaviour w, 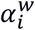 is the effect of treatment i, and W is the reference feeding behaviour. Submodel (2) was compared to a model were the differences between the observations were only due to the plates, whatever the treatment

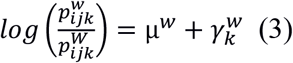

Models (2) and (3) were compared using the Akaike Information Criterion (AIC) [47]. In the two experiments, model (2) was chosen by the AIC criterion, meaning that probabilities that one cage falls in one given behaviour group rather than another depends more on the treatment (product concentration) than the plate it belonged to.

The multinomial regression (2) provided an estimation of the probabilities associated with each treatment, using

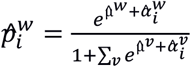

that were used for the graphical representations (Fig 4).

A Wald test was performed to compare the treatments [48]. Contrasts between two treatments i and i’ are computed as the differences 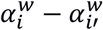. A positive value of the contrast means that the proportion of feeding type w relative to feeding type W is greater in treatment i than in treatment i’. With multinomial regression, the choice of the reference is tricky. The best reference is the category were the observations are equally distributed between the treatments. We chose the feeding type F as the reference for the quinine experiment, and the feeding behaviour E for the NeemAzal experiment.

## Acknowledgments

This work was supported by a scholarship from the Islamic Bank of Development to Inoussa Sanane (N° BID: 600033174) and a grant from the BASC labex (ITEMAIZE project) supported by IDEX Paris-Saclay. We are grateful to Stephane Dallongeville (Institut Pasteur), for his constant help and support for the development of the ICY plugins. We warmly thank members of J Legrand’s and F Marion-Poll’s laboratories for discussion and support during this work and R Jeannette for his help for the insects rearing.

## Supplementary information

**SI-Table 1:**
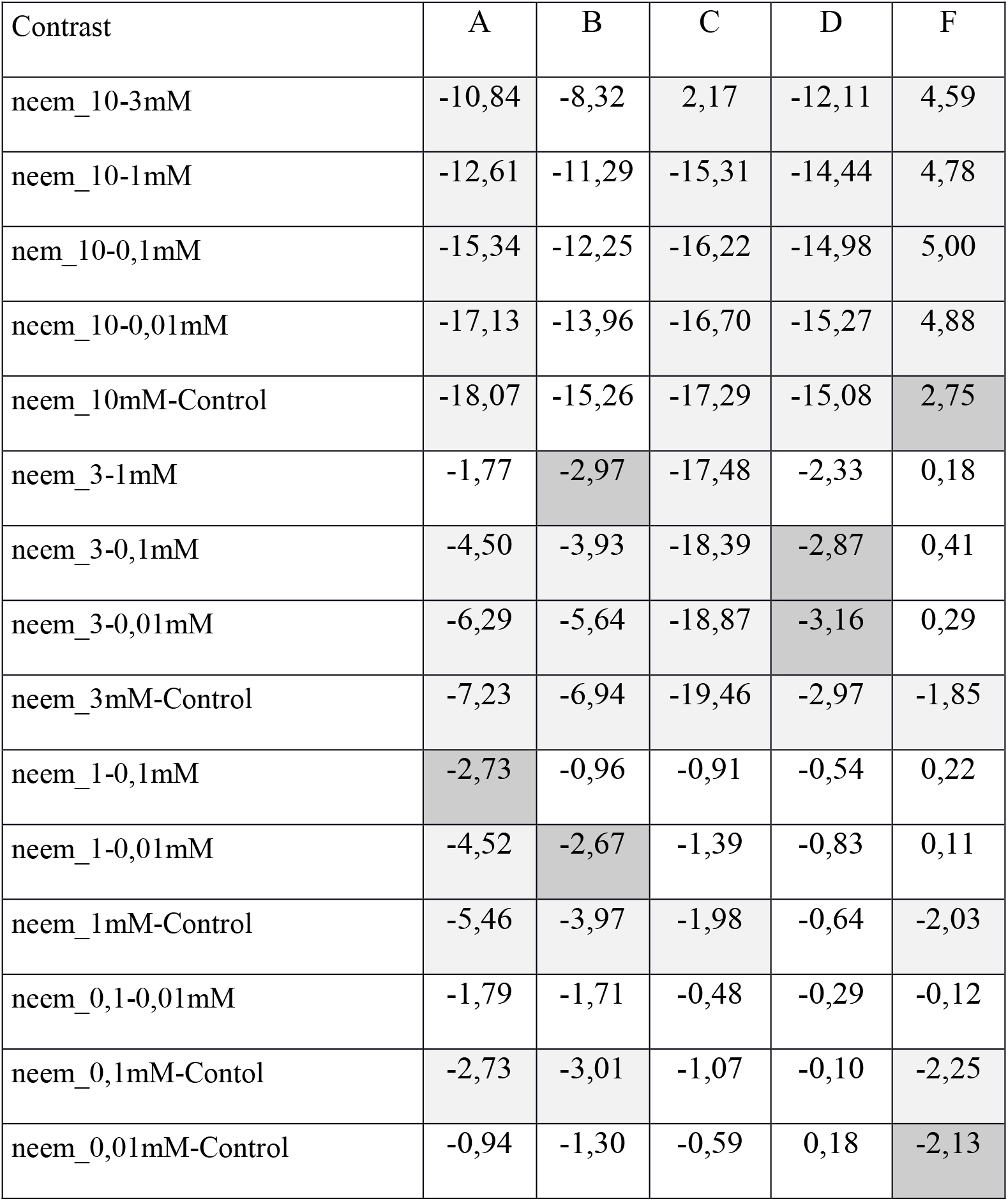
Contrasts between the different concentrations of NeemAzal treatments within each typology of feeding behaviour. Contrasts are expressed on the logit scale, relative to the reference feeding behavioural category E. For each contrast and each behavioural type, the levels of significance is indicated by the colour filling. Darkgray = significant at the 1%o risk level.1. Lightgrey = significant at the 1% risk level. White = non-significant at the 1% risk level.

| Contrast | A | B | C | D | F |
| --- | --- | --- | --- | --- | --- |
| neem_10-3mM | -10,84 | -8,32 | 2,17 | -12,11 | 4,59 |
| neem_10-1mM | -12,61 | -11,29 | -15,31 | -14,44 | 4,78 |
| neem_10-0,1mM | -15,34 | -12,25 | -16,22 | -14,98 | 5,00 |
| neem_10-0,01mM | -17,13 | -13,96 | -16,70 | -15,27 | 4,88 |
| neem_10mM-Control | -18,07 | -15,26 | -17,29 | -15,08 | 2,75 |
| neem_3-1mM | -1,77 | -2,97 | -17,48 | -2,33 | 0,18 |
| neem_3-0,1mM | -4,50 | -3,93 | -18,39 | -2,87 | 0,41 |
| neem_3-0,01mM | -6,29 | -5,64 | -18,87 | -3,16 | 0,29 |
| neem_3mM-Control | -7,23 | -6,94 | -19,46 | -2,97 | -1,85 |
| neem_1-0,1mM | -2,73 | -0,96 | -0,91 | -0,54 | 0,22 |
| neem_1-0,01mM | -4,52 | -2,67 | -1,39 | -0,83 | 0,11 |
| neem_1mM-Control | -5,46 | -3,97 | -1,98 | -0,64 | -2,03 |
| neem_0,1-0,01mM | -1,79 | -1,71 | -0,48 | -0,29 | -0,12 |
| neem_0,1mM-Control | -2,73 | -3,01 | -1,07 | -0,10 | -2,25 |
| neem_0,01mM-Control | -0,94 | -1,30 | -0,59 | 0,18 | -2,13 |

**SI-Table 2:** Contrasts between the different concentrations of Quinine treatments within each typology of feeding behaviour. Contrasts are expressed on the logit scale, relative to the reference behavioural type F. For each contrast and each behavioural type, the levels of significance is indicated by the colour filling. Darkgray = significant at the 1%o risk level. Lightgrey = significant at the 1% risk level. White = non-significant at the 1% risk level.

|  | A | B | C | D | E |
| --- | --- | --- | --- | --- | --- |
| quin_100-10mM | -1,3 | -0,08 | 16,06 | 16,47 | 15,12 |
| quin_100-1mM | 0,14 | 0,49 | 0,742 | 0,657 | 1,099 |
| quin_100-0,1mM | -1,94 | -0,75 | -0,42 | 1,216 | 0,118 |
| quin_100-0,01mM | 0,05 | 0,41 | 1,738 | 2,683 | 15,17 |
| quin_100mM-Control | -0,62 | 0,55 | -0,13 | 1,453 | 1,558 |
| quin_10-1mM | 1,44 | 0,57 | -15,3 | -15,8 | -14 |
| quin_10-0,1mM | -0,64 | -0,67 | -16,5 | -15,3 | -15 |
| quin_10-0,01mM | 1,35 | 0,49 | -14,3 | -13,8 | 0,054 |
| quin_10mM-Control | 0,68 | 0,63 | -16,2 | -15 | -13,6 |
| quin_1-0,1mM | -2,08 | -1,24 | -1,16 | 0,559 | -0,98 |
| quin_1-0,01mM | -0,1 | -0,09 | 0,996 | 2,026 | 14,07 |
| quin_1mM-Control | -0,76 | 0,05 | -0,88 | 0,796 | 0,46 |
| quin_0,1-0,01mM | 1,98 | 1,15 | 2,16 | 1,467 | 15,05 |
| quin_0,1mM-Control | 1,32 | 1,29 | 0,288 | 0,237 | 1,441 |
| quin_0,01mM-Control | -0,66 | 0,14 | -1,87 | -1,23 | -13,6 |

